# Linking spatial self-organization to community assembly and biodiversity

**DOI:** 10.1101/2021.06.20.449156

**Authors:** Bidesh K. Bera, Omer Tzuk, Jamie J. R. Bennett, Ehud Meron

**Author notes:** **For correspondence** (EM). Department of Industrial Engineering Faculty of Engineering Tel-Aviv University, Tel Aviv 6997801, Israel (Omer Tzuk).

## Abstract

Drier climates impose environmental stresses on plant communities that may result in community reassembly and threatened ecosystem services, but also may trigger self-organization in spatial patterns of biota and resources, which act to relax these stresses. The complex relationships between these counteracting processes – community reassembly and spatial self-organization – have hardly been studied. Using a spatio-temporal model of dryland plant communities and a trait-based approach, we study the response of such communities to imposed water stress of increasing degrees. We first show that spatial patterning acts to reverse shifts from fast-growing species to stress-tolerant species, as well as to reverse functional-diversity loss. We then show that spatial re-patterning buffers the impact of further stress on community structure. Finally, we identify multistability ranges of uniform and patterned community states and use them to propose forms of non-uniform ecosystem management that integrate the need for provisioning ecosystem services with the need to preserve community structure.

## Introduction

The structure of plant communities – their composition and diversity – forms the foundation of many ecosystem services on which human well-being crucially depends. These include provisioning services such as food, fresh water, wood and fiber; regulating services such as flood regulation and water purification; cultural services such as recreation and aesthetic enjoyment; and supporting services such as soil formation, photosynthesis, and nutrient cycling (***Duraiappah and Naeem, 2005***). These services are at risk due to potential changes in the composition and diversity of plant communities as a result of global warming and the development of drier climates (***Harrison et al., 2020***; ***Grünzweig and et al., 2021***). Understanding the factors that affect community structure in varying environments calls for integrated studies of mechanisms operating at different levels of organization, from phenotypic changes at the organism level, through *intra*specific interactions at the population level, to *inter*specific interactions at the community level (***Gratani, 2014***; ***Falik et al., 2003***; ***Bertness and Callaway, 1994***; ***Pérez-Ramos et al., 2019***). Of these mechanisms, the role of intraspecific interactions in driving community dynamics through spatial self-organization, has hardly been studied (***Vandermeer and Yitbarek, 2012***; ***Bonanomi et al., 2014***; ***Cornacchia et al., 2018***; ***Zhao et al., 2019***; ***O’Sullivan et al., 2019***).

Spatial self-organization in periodic and non-periodic patterns of biota and resource, driven by intraspecific competition that leads to partial plant mortality, is widely observed in stressed environments (***Rietkerk and van de Koppel, 2008***). An important class of these phenomena are vegetation patterns in drylands. In sloped terrains these patterns commonly appear as parallel vegetation stripes oriented perpendicular to the slope direction (***Lefever and Lejeune, 1997***; ***Valentin et al., 1999***; ***Bastiaansen et al., 2018***). In flat terrains, with no preferred direction imposed by slope or wind, stripe-like configurations often appear as labyrinthine patterns. However, in such terrains two additional pattern morphologies are often observed; nearly periodic patterns of bare-soil gaps in otherwise uniform vegetation, and nearly periodic patterns of vegetation spots in otherwise uniform bare soil (***Rietkerk et al., 2004***; ***Deblauwe et al., 2008***; ***Borgogno et al., 2009***; ***Meron, 2018***). Spatial self-organization may not necessarily result in periodic patterns; according to pattern-formation theory (***Meron, 2015***; ***Knobloch, 2015***) it can also result in non-periodic patterns, such as single or randomly scattered vegetation spots in otherwise bare soil, randomly scattered bare-soil gaps in otherwise uniform vegetation and others (***Tlidi et al., 2008***; ***Dawes and Williams, 2015***; ***Parra-Rivas and Fernandez-Oto, 2020***; ***Jaïbi et al., 2020***; ***Zelnik et al., 2015***). The driving mechanisms of these self-organized vegetation patterns are scale-dependent positive feedback loops between local vegetation growth and water transport toward the growth location (***Rietkerk and van de Koppel, 2008***; ***Meron, 2019***).

Vegetation patterns involve not only spatial distributions of plant biomass, but also less-visible distributions of soil-water, nutrients, soil biota, and possibly toxic substances (***Paz-Kagan et al., 2019***; ***Inderjit. and Duke, 2003***; ***De Deyn et al., 2004***; ***van der Putten et al., 2013***). The various habitats that these self-organizing distributions form lead to niche differentiation and community reassembly (***Weiher et al., 2011***). Thus, spatial self-organization and community dynamics are intimately-coupled processes that control community composition and diversity. Understanding this unexplored coupling is essential for assessing the impact of global warming and drier climates on community structure and ecosystem services.

In this paper we incorporate spatial self-organization into community-assembly studies, using a mathematical model of dryland plant communities. Our model study provides three new insights, illustrated in Fig. 1: (i) Spatial self-organization acts to reverse community-structure changes induced by environmental stress, (ii) it buffers the impact of further stress, and (iii) it offers new directions of ecosystem management that integrate the need for provisioning ecosystem services with the need to conserve community structure. More specifically, using a trait-based approach (***Diaz and Cabido, 2001***), we show that drier climates shift the composition of spatially uniform communities towards stress-tolerant species, and reduce functional diversity. By contrast, self-organization in spatial patterns, triggered by these droughts, shift the composition back to less tolerant species that favor investment in growth, and increase functional diversity. Once patterns are formed, spatial self-organization provides various pathways to relax further stresses without significant changes in community composition and diversity. Furthermore, multistability ranges of uniform and patterned states open up new opportunities for grazing and foddering management by forming mixed community states of increased functional diversity.

**Figure 1.**
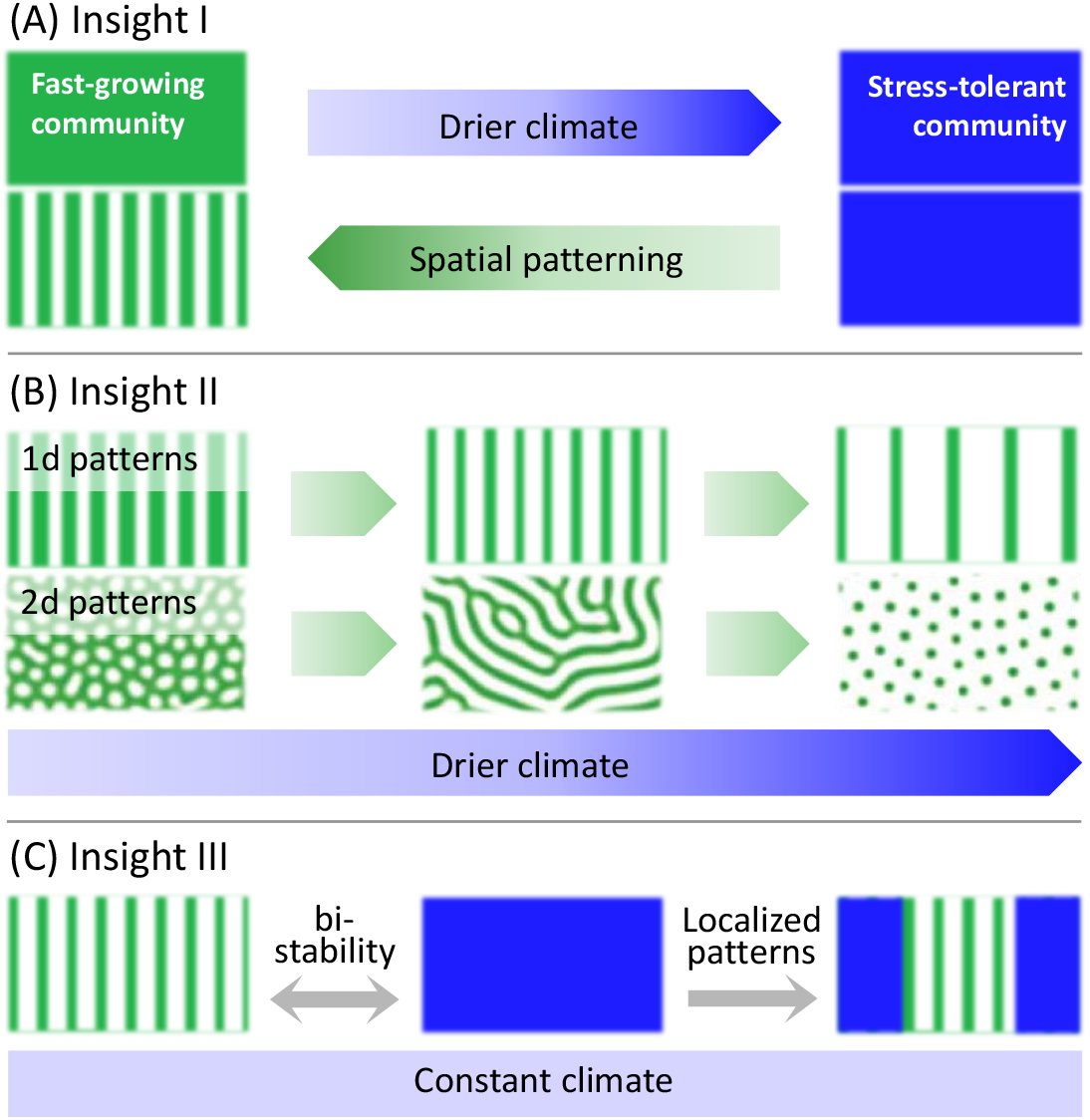
A schematic illustration of the three insights that the model study provides. (A) Insight I: A drier climate shifts an original spatially uniform community of fast-growing plants, denoted by a green color, to a uniform community of stress-tolerant plants, denoted by blue color. Spatial patterning induced by the drier climate shifts the community back to fast growing plants. (B) Insight II: Once patterns have formed a yet drier climate has little effect on community structure – all patterned community states consist of fast-growing plants (green). This is because of further processes of spatial self-organization that increase the proportion of water-contributing bare-soil areas and compensate for the reduced precipitation. In 1d patterns, these processes involve thinning of vegetation patches or transitions to longer wavelength patterns. In 2d patterns, these processes involve morphological transitions from gap to stripe patterns and from stripe to spot patterns. (C) Insight III: Localized patterns in a bistability range of uniform and patterned community states significantly increase functional diversity, as they consist of both stress-tolerant (blue) and fast-growing (green) species. Such patterns can be formed by nonuniform biomass removal as an integral part of a provisioning ecosystem service.

## Results

### A model for spatial assembly of dryland plant communities

The model we study largely builds from foundations introduced earlier (***Meron, 2016***). These foundations capture three pattern-formation mechanisms associated with three forms of water transport: overland water flow, soil-water diffusion, and water conduction by laterally spread roots (***Meron, 2019***). In this study we focus, for simplicity, on a single mechanism associated with overland water flow. The mechanism induces a stationary nonuniform (Turing) instability of uniform vegetation, leading to periodic vegetation patterns. The instability can be understood in terms of a positive feedback loop between local vegetation growth and overland water flow towards the growth location: an area with an incidental denser vegetation draws more water from its surrounding areas than the latter do, which accelerates the growth in that area and decelerates the growth in the surrounding areas, thus amplifying the initial nonuniform perturbation. This feedback loop is also called a scale-dependent feedback because of the positive effect on vegetation growth at short distances and negative effect at longer distances (***Rietkerk and van de Koppel, 2008***). The reason why an area of denser vegetation draws more water from its surroundings is rooted in the differential infiltration of overland water into the soil that develops – high in denser vegetation and low in sparser vegetation, as illustrated in Fig. 2. Several factors contribute to this process, including denser roots in denser-vegetation areas, which make the soil more porous, and lower coverage of the ground surface in denser-vegetation areas by physical or biological soil crusts, which act to reduce the infiltration rate. The differential infiltration induces overland water flow towards areas of denser vegetation that act as sinks.

**Figure 2.**
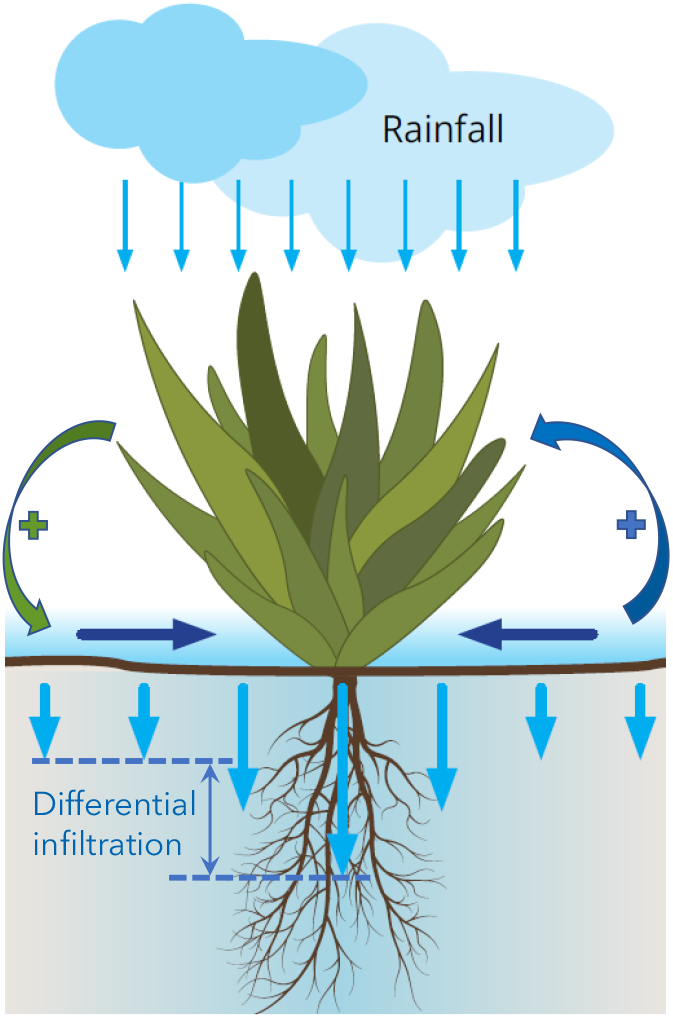
Illustration of overland water flow towards vegetation patches (horizontal arrows), induced by differential infiltration: low in bare soil (short vertical arrows) and high in vegetation patches (long arrows). The round arrows denote the positive feedback loop between vegetation growth and overland water flow towards the growth location. This feedback loop destabilizes uniform vegetation to form vegetation patterns, and acts to stabilize these patterns once formed.

We quantify the plant community by introducing a dimensionless trait parameter (***Nathan et al., 2016***; ***Tzuk et al., 2019***; ***Yizhaq et al., 2020***), *χ* ∈ (0, 1], that represents a tradeoff between plant investment in shoot growth vs. investment in tolerance to water stress, so that *χ* → 0 represents plants that invest mostly in growth, while *χ* → 1 represents plants that invest mostly in tolerating water stress. Using this parameter, the pool of species is divided into *N ≫* 1 functional groups, where all species within the *i*th group have *χ* values in the small interval Δ_*χ*_ = 1/*N* that precedes *χ*_*i*_ = *i*Δ_*χ*_. Spatial self-organization is captured by including a pattern-forming feedback associated with overland water flow towards areas of denser vegetation where the infiltration rate is higher (***Meron, 2019***). Altogether the model consists of the following system of PDEs for biomass variables, *B*_*i*_ (*i* = 1, …, *N*), representing the areal biomass densities of the *N* functional groups, and two water variables, *W* and *H*, representing below-ground and above-ground water, respectively, all in units of kg/m^2^:

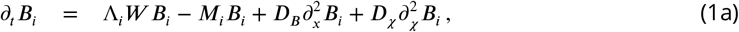

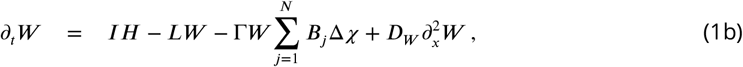

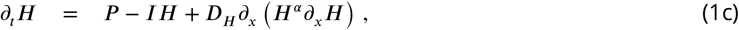

where the second ‘trait derivative’, 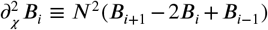, represents mutations at a very small rate *D*_*χ*_. In these equations the growth rate of the *i*th functional group, Λ_*i*_, the infiltration rate of above-ground water into the soil, *I*, and the evaporation rate, *L*, are given by the expressions:

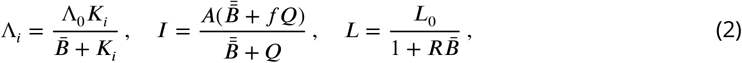

where 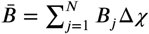 and 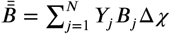.

The biomass dependence of the growth rates, Λ_*i*_, models competition for light and accounts for growth attenuation due to shading. That attenuation is quantified by the parameters *K*_*i*_; high *K*_*i*_ values represent plant species investing preferably in shoot growth that are less affected by shading. Note that the growth rate also includes attenuation due to self-shading (***Noy-Meir, 1975***). The parameter Λ_0_ represents the growth rate at low biomass values for which competition for light is absent. The biomass dependence of the infiltration rate, *I*, is responsible for differential infiltration, quantified by the dimensionless parameter 0 ≤ *f ≤*1 and the parameter *Q*. Low *f* values represent highly differential infiltration, low in bare soil and high (*Q*-dependent) in vegetation patches, and constitute an important element in the pattern-forming feedback associated with overland water flow towards denser vegetation patches (***Meron, 2019***). The biomass dependence of the evaporation rate, *L*, accounts for reduced evaporation in vegetation patches due to shading, quantified by the parameter *R*. The parameter *L*_0_ represents evaporate rate in bare soil. Tolerance to water stress is modeled through the mortality parameters *M*_*i*_ – a plant investment in tolerating stress reduces the mortality rate. For simplicity, we choose to describe here overland water flow as a linear diffusion problem by setting *α* = 0. Although that process is nonlinear, the qualitative results and conclusions reported here do not depend on that choice (see ***Gilad et al***. (***2004***) for the choice *α* = 1). Additional model parameters are *P*, representing mean annual precipitation, Γ, representing the rate of water uptake by plants’ roots, *D*_*B*_ representing seed dispersal rate, *D*_*W*_ quantifying lateral soil-water diffusion, and *D*_*H*_ quantifying overland water flow.

As pointed out earlier, we distinguish between different species through the different tradeoffs they make between shoot growth and tolerance to water stress. We capture this tradeoff using the parameters *K*_*i*_ and *M*_*i*_ through the tradeoff relations:

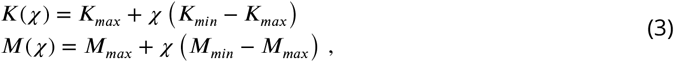

where *K*_*i*_ = *K*(*χ*_*i*_) and *M*_*i*_ = *M*(*χ*_*i*_). According to these relations, *χ* → 0 represents the functional group (*K*_*max*_, *M*_*max*_) with highest investment in growth and lowest investment in tolerance (highest mortality), while *χ* = 1 represents the functional group (*K*_*min*_, *M*_*min*_) with lowest investment in growth and highest investment in tolerance. This tradeoff is likely to affect the contributions, *Y*_*i*_, of the various functional groups to the infiltration rate *I*; denser roots associated with lower-*χ* species (increased investment in shoot growth) make the soil more porous and increase the infiltration rate. An additional contribution to that effect is lower soil-crust coverage in patches of lower-*χ* species. We therefore assume the following form for *Y*_*i*_:

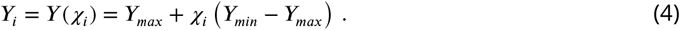

We use numerical continuation methods (AUTO (***Doedel, 1981***)) to study the model equations for single functional groups, and numerical time-integration in the composed trait-space plane to study the full community model. We use periodic boundary conditions in *x* and zero-flux conditions in *χ*. Initial conditions are chosen to contain all functional groups, even if at diminishingly small biomass values. Such small values represent the presence of seeds that remain viable even when they cannot germinate (***DeMalach et al., 2021***). Throughout the paper we use the following parameter values: *N* = 128, Λ_0_ = 0.032 m^2^/(kg y), Γ = 20.0 m^2^/(kg y),*A*= 40.0 y^−1^, *R* = 10.0 m^2^/kg, *L*_0_ = 4.0 y^−1^, *f* = 0.01, *Q* = 0.06 kg/m^2^, *K*_*min*_ = 0.1 y^−1^, *K*_*max*_ = 0.6 kg/m^2^, *M*_*min*_ = 0.5 y^−1^, *M*_*max*_ = 0.9 y^−1^, *Y*_*min*_ = 0.5, *Y*_*max*_ = 1.5, *D*_*B*_ = 1.0 m^2^/y, *D*_*W*_ = 10^2^ m^2^/y, *D*_*H*_ = 10^4^ m^2^/y, *D*_*χ*_ = 10^−6^ y^−1^. Values of other parameters are as stated in the following.

We wish to point out that the results presented here are not sensitive to this particular choice of parameter values, and similar results have been obtained with other sets of parameter values. The main constraint on this choice is the need to capture the Turing instability of the uniform vegetation state, and the need to define the tradeoff relations such that the Turing threshold split the community into functional groups that form and do not form periodic patterns. Since vegetation patterns are observed on spatial scales that differ by orders of magnitude, from periodicity of tens of centimeters for herbaceous vegetation to periodicity of tens of meters for woody vegetation (***Rietkerk et al., 2004***), we focus on generic community aspects associated with vegetation patterning, rather than attempt to model a particular ecosystem with a specific spatial scale.

### Single functional-group states

It is instructive to consider first solutions of a model for a single functional group, *χ* = 1. As the bifurcation diagram in Fig. 3a shows, a uniform vegetation solution (*UV*) exists and is stable at sufficiently high precipitation (*P*), but loses stability in a subcritical Turing bifurcation (***Meron, 2018***) as *P* is decreased below a threshold *P*_*T*_ (*χ*). That instability creates a bistability range of uniform vegetation and periodic patterns (*PP*) where hybrid states (*HS*), consisting of patterned domains of increasing size in otherwise uniform vegetation, exist (***Knobloch, 2015***; ***Zelnik et al., 2015***). Besides the periodic-pattern solution that appears at the Turing bifurcation (*P P*_*W L*=81_), many more periodic solutions appear, as *P* further decreases, with longer wavelengths (WL) (***Zelnik et al., 2013***; ***Siteur et al., 2014***). The second periodic solution shown in the diagram (*P P*_*W L*=150_) has a wavelength almost twice as long. A solution describing bare soil (*BS*), devoid of vegetation, exists at all *P* values but is stable only below a threshold value *P*_*B*_. Similar bifurcation diagrams are obtained for lower *χ* values, but the existence and stability ranges of the various solutions change. As Fig. 3b shows, the uniform-vegetation state of species investing mostly in growth (low *χ*) lose stability to periodic patterns at higher *P* values. Also, the bare-soil state remains stable at higher *P* values. These results imply that species investing in growth have a stronger tendency to form patchy vegetation, and are more at risk of mortality (collapse to bare soil) as a result of disturbances.

**Figure 3.**
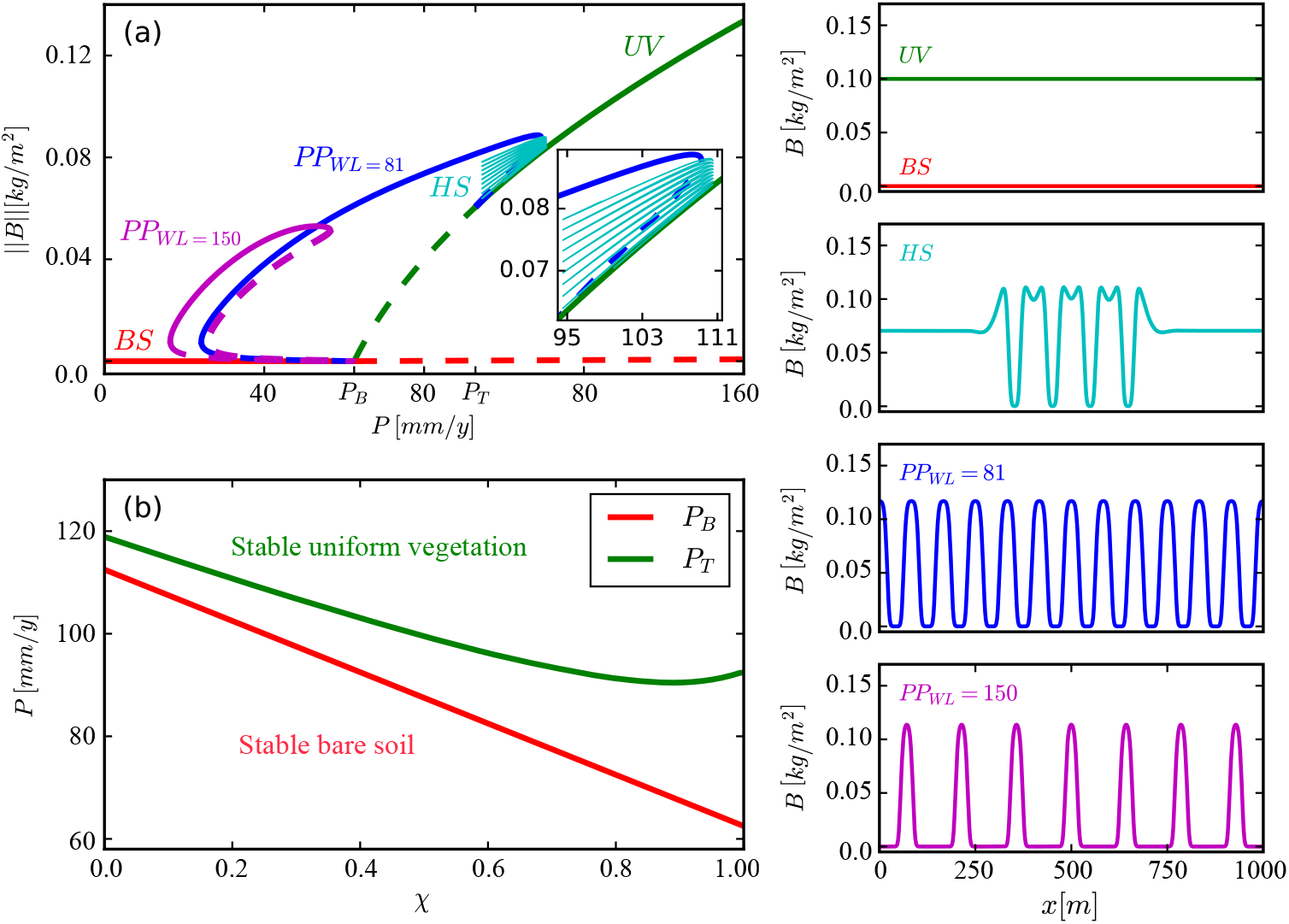
Existence and stability ranges of various solutions of a model for a single functional group. (a) A bifurcation diagram showing the *L*2-norm of the biomass density vs. precipitation for *χ* = 1. The colors and corresponding labels denote the different solution branches: uniform vegetation (*UV*), periodic patterns at different wavelengths (*PP*), hybrid states consisting of pattern domains in otherwise uniform vegetation (*HS*), and bare soil (*BS*). Solid (dashed) lines represent stable (unstable) solutions. Example of spatial profiles of these solutions are shown in the panels on the right. (b) Instability thresholds of uniform vegetation, *P*_*T*_, and of bare soil, *P*_*B*_, as functions of *χ*.

### Effects of spatial patterning on community assembly

What forms of community assemblages can emerge when the *N* functional groups are allowed to interact and compete with one another? Figure 4 shows the assembly of a spatially uniform community, where all functional groups asymptotically form uniform vegetation (*P > P*_*T*_ (*χ* → 0), see Fig. 3b). Because of species competition for water and light, a particular community assemblage develops, often characterized by a hump-shape biomass distribution with a most abundant group (maximal biomass) at *χ*_*max*_. This distribution contains information about two community-level properties of interest here: the community’s composition, quantified by *χ*_*max*_, and its functional diversity. The latter represents the diversity of functional traits around *χ*_*max*_ (***Diaz and Cabido, 2001***).

**Figure 4.**
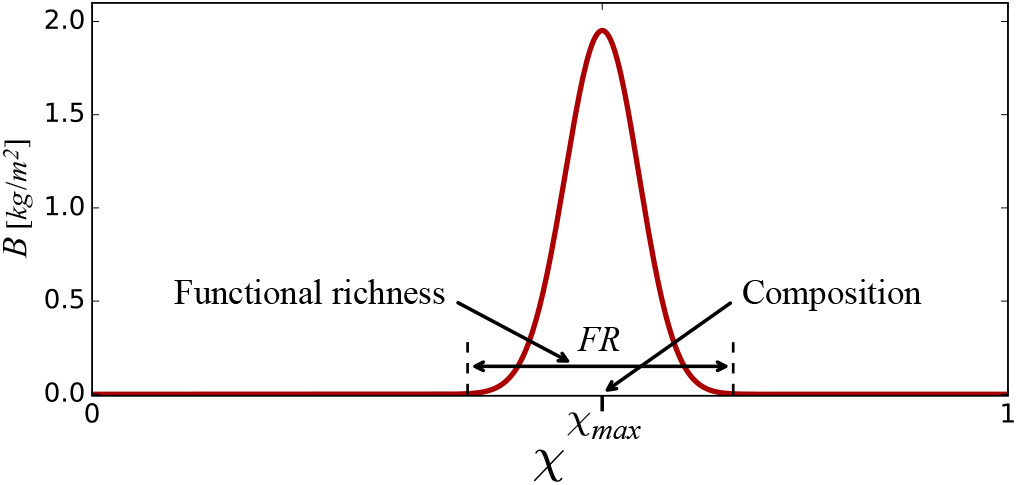
Community-level properties derivable from model solutions. The biomass distribution along the trait axis *χ* contains information about several community-level properties, including community composition, quantified by the position, *χ*_*max*_, of the most abundant functional group, and functional richness, quantified by the distribution width, *FR*, at diminishingly small biomass values.

We measure functional diversity using two metrics, functional richness, *FR*, and functional evenness, *FE* (***Mason et al., 2005***). The first metric is given by the extent of the biomass distribution around *χ*_*max*_, as Fig. 4 illustrates. The second metric contains information about the abundance of functional groups in the community and how even the abundance is among the groups. We use here the following analogue of the Shannon diversity index,

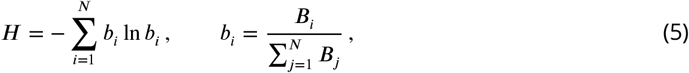

and the related index of Pielou for functional evenness (***Pielou, 1966***),

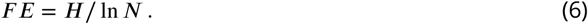

How do plant communities respond to progressively drier climates, mimicked here by precipitation downshifts of increasing strength? Figure 5 shows asymptotic biomass distributions for decreasing precipitation values. At *P*_1_ = 150mm/y (panel a), a spatially uniform hump-shape community develops, characterized by a symmetric distribution of functional groups around a most abundant group at *χ*_*max*_ = *χ*_0_ = 0.62, and by functional richness *FR*_0_ = 0.29. Lowering the precipitation to *P*_2_ = 100mm/y (panel b), results in a spatially uniform community shifted to species that better tolerate water stress (higher *χ*), now distributed around a most abundant function group at *χ*_0_ = 0.78, and reduced functional richness, *FR*_0_ = 0.25.

**Figure 5.**
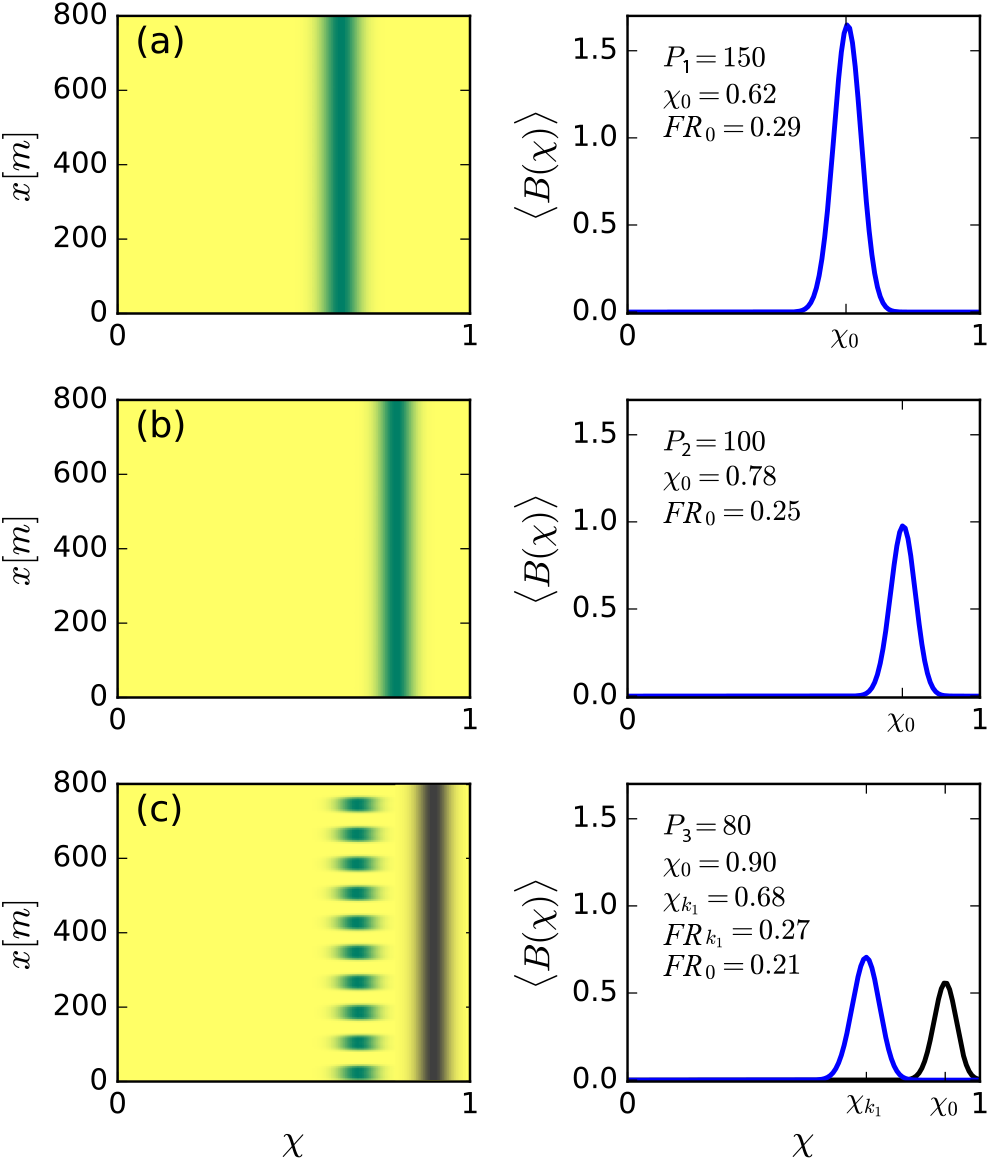
Community reassembly in response to precipitation downshifts. Left panels show biomass distributions in the trait (*χ*) – space (*x*) plane for the specified precipitation rates *P*_1_, *P*_2_, *P*_3_. Right panels show biomass profiles along the *χ* axis averaged over space. (a,b) A precipitation downshift from *P* = 150 mm/y to 100 mm/y, starting with a uniform community, results in a uniform community shifted to more tolerant species (higher *χ*), and of lower functional richness (*FR*). (c) Further decrease to *P* = 80 mm/y results in a patterned community that is shifted back to species investing in growth (lower *χ*), and has higher functional richness. The biomass distributions in black refer to the unstable uniform community.

Lowering the precipitation further yet to a value, *P*_3_ = 80mm/y, below the Turing threshold *P*_*T*_, results in spatial patterning (panel c). Interestingly, the community composition is now shifted back to species that favor investment in growth (lower *χ*), and the functional richness increases rather than continue to decrease; the patterned community is distributed around 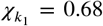 and its functional richness is 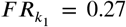. Panel (c) also shows (in black) the biomass distribution of the unstable spatially-uniform community at *P*_3_ = 80mm/y, which continues the trend of panels (a,b) and demonstrates the significant change in community structure that spatial self-organization induces.

Once periodic patterns form, a further decrease in precipitation does not result in significant community-structure changes, unlike the case of uniform communities, as Fig. 6 shows. While spatially uniform communities move to higher *χ* values with decreasing precipitation, as the monotonically decreasing graph *χ*_*max*_ = *χ*_0_(*P*) shows, spatially patterned communities remain largely unchanged, as the nearly horizontal graphs 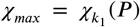 and 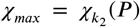 show. The first graph represents the periodic pattern that emerges at the Turing instability point *P*_*T*_. Along this graph, as *P* is reduced, the number of vegetation patches remains constant, but their size (along the *x* axis) significantly reduces (compare the insets at *P* = *P*_3_ and *P* = *P*_4_ in Fig. 6). Furthermore, the patches span the same range of functional groups (patch extension along the *χ* axis), i.e. retain their functional richness, and their most abundant functional group, 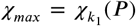, does not change. Thus, as *P* is reduced, the patterned community that emerges at the Turing instability hardly changes in terms of pattern wavenumber *k*_1_ or wavelength 2*π*/*k*_1_, and in terms of community structure, but the abundance of all functional groups reduces significantly as the vegetation patches become thinner along the *x* axis.

**Figure 6.**
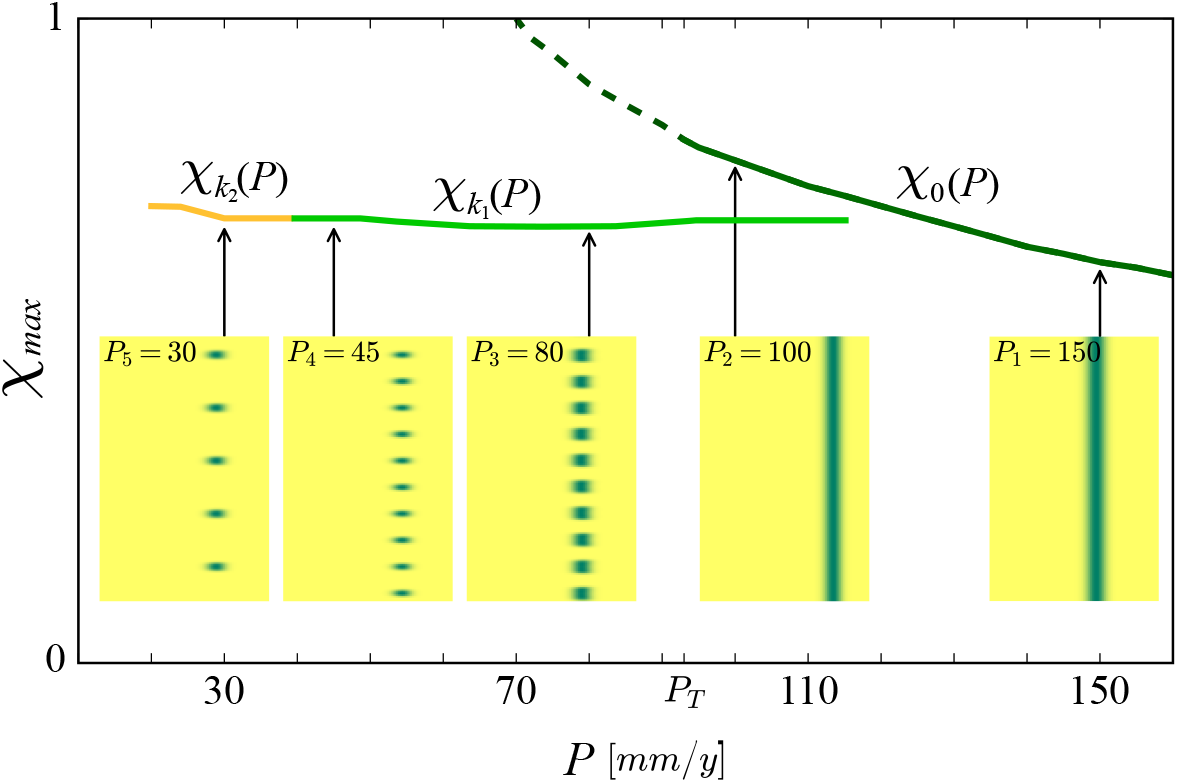
The buffering effect of spatial patterning on community structure. Shown is a partial bifurcation diagram depicting different forms of community assembly along the precipitation axis, as computed by integrating the model equations in time. Stable, spatially uniform communities, *χ*_0_(*P*) (solid dark-green line), shift to stress-tolerant species (higher *χ*), as precipitation decreases. When the Turing threshold, *P*_*T*_, is traversed, spatial self-organization shifts the community back to growth species (lower *χ*), and keep it almost unaffected as the fairly horizontal solution branches, 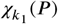 (light green line), representing periodic patterns of wavenumber *k*_1_, and 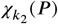 (yellow line), representing patterns of lower wavenumber *k*_2_, indicate. The insets show biomass distributions in the (*χ, x*) plane for representative precipitation values. The unstable solution branch describing uniform vegetation (dashed line) was calculated by time integration of the spatially-decoupled model.

The second graph 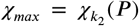 represents a periodic vegetation pattern consisting of fewer patches (along the *x* axis). Their extension along the *χ* axis remains approximately constant (functional richness hardly changes), and the same holds for the most abundant functional group, 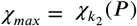. Thus, the effect of further precipitation decrease is a transition to periodic pattern of longer wavelength and reduced abundance, but the community structure (composition and richness) remains almost unaffected.

### Effects of multistability on community assembly

The discussion so far focused on spatial self-organization in periodic patterns. However, the bifurcation diagram of Fig. 3, obtained for a single functional group, suggests the possible existence of non-periodic or disordered patterns as well, associated with (i) homoclinic snaking (***Knobloch, 2015***) in the bistability range of uniform and periodic vegetation, which gives rise to a multitude of additional non-periodic hybrid states (***Meron, 2019***), (ii) multiplicity of stable periodic patterns of different wavenumbers within the Busse balloon (***Sherratt, 2013***; ***Zelnik et al., 2013***; ***Siteur et al., 2014***; ***Bastiaansen et al., 2018***). We focus here on the first form of multistability and show in Fig. 7 three examples of hybrid states that differ in the size of the patterned domain relative to the uniform domain. These model solutions were obtained by starting from similar hybrid solutions of a single functional-group model as initial conditions (Fig. 3). The community dynamics that develop in the course of time leads to different communities in the two domain types; stress-tolerant species in the uniform domains, and fast growing species in the patterned domains. As a consequence, the global, whole-system functional richness increases significantly, compared to the richness associated with a purely uniform state or a purely patterned state, as the right panels in Fig. 7 indicate (compare with Fig. 5b). Furthermore, the relative size of the patterned domains affects the functional evenness, *FE* (see Eq. 6). A relatively high *FE* value is obtained for uniform and patterned domains of comparable sizes (Fig. 7b) and low evenness when the relative size of the patterned domain is either small or large (Fig. 7a,c).

**Figure 7.**
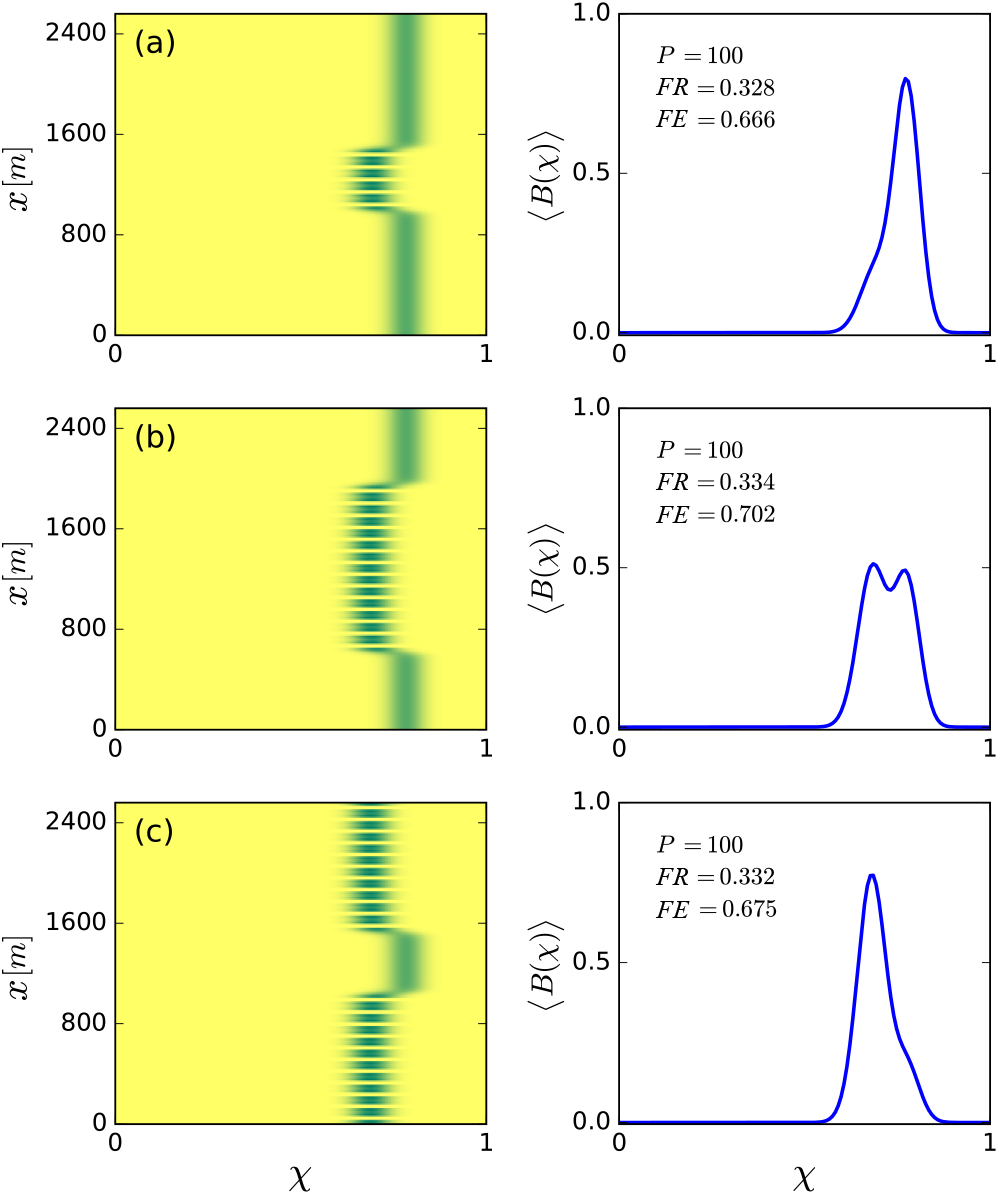
Increased functional diversity of hybrid states and evenness control. Left panels show biomass distributions of different hybrid states in the trait (*χ*) – space (*x*) plane. Right panels show biomass profiles along the *χ* axis averaged over space. The functional richness, *FR*, of all hybrid states is almost equal and higher than that of purely uniform or purely patterned states (compare with panel b in Fig. 5), but their functional evenness, *FE*, differs – high for patterned and uniform domains of comparable sizes (b) and low for small (a) and large (c) pattern-domain sizes. Calculated for a precipitation rate *P* = *P*_2_ = 100 mm/y.

## Discussion and conclusion

The results described above provide three insights into the intimate relationships between spatial self-organization, community assembly and ecosystem management, as illustrated in Fig. 1 and explained below.

### Insight I: Spatial patterning acts to reverse community-structure changes induced by environmental stress

According to Fig. 5, reduced precipitation shifts spatially uniform communities to stress-tolerant species (higher *χ* values), but when the Turing threshold is traversed and self-organization in periodic spatial patterns occurs, this trend is reversed and a shift back to species investing in growth takes place. This surprising change of community structure reflects the complex nature of ecosystem response to varying environments, which can employ mechanisms operating in parallel at different organization levels. The composition shift to higher *χ* values, as *P* decreases but still remains above the Turing threshold *P*_*T*_, is driven by community-level processes, whereby *inter*specific competition results in a community consisting of species that are better adapted to water stress, and of lower functional richness. By contrast, the composition shift to lower *χ* values, once the Turing threshold is traversed, is driven by population-level processes of spatial self-organization, whereby *intra*specific competition results in partial mortality and the appearance of bare-soil patches. These patches provide an additional source of water to adjacent vegetation patches, besides direct rainfall, through overland water flow (***Meron, 2019***). That additional resource compensates for the reduced precipitation and relaxes the local water stress at vegetation patches. The resulting ameliorated growth conditions favor species investing in growth (lower *χ*), and increase functional richness.

### Insight II: Spatial re-patterning buffers community-structure changes

Once periodic patterns form, a further decrease of precipitation does not result in significant community-structure changes, as Fig. 6 shows. This is because of additional forms of spatial self-organization that buffer the impact of decreasing precipitation. The first of which is partial plant mortality that results in vegetation patches of reduced size (compare the insets at *P* = *P*_3_ and *P* = *P*_4_ in Fig. 6). These patches benefit from increased water availability due to the larger water-contributing bare-soil patches that surround them. This response form does not involve a change in the number of patches or pattern’s wavenumber, and occurs along the branch of any periodic solution, including those shown in Fig. 6, 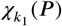 and 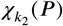. A second form of spatial self-organization involves plant mortality that results in patch elimination and wavenumber reduction (***Yizhaq et al., 2005***; ***Siteur et al., 2014***), such as the transition from 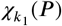 to 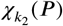 in Fig. 6 (see insets at *P* = *P*_4_ and *P* = *P*_5_). Like in the first form, any remaining vegetation patch benefits from larger bare-soil patches surrounding it and, thus, from higher water availability.

These forms of spatial self-organization apply also to two-dimensional systems, especially to gently sloped terrains, where quasi-one-dimensional patterns of stripes oriented perpendicular to the slope direction occupy wide precipitation ranges (***Deblauwe et al., 2010***) and are widely observed in nature (***Valentin et al., 1999***; ***Deblauwe et al., 2012***; ***Bastiaansen et al., 2018***). Two-dimensional systems, however, allow for additional forms of patterning and re-patterning, which we have not studied in this work (***von Hardenberg et al., 2001***; ***Rietkerk et al., 2002***; ***Lejeune et al., 2004***; ***Gowda et al., 2014***; ***Meron, 2019***). In flat terrains uniform vegetation responds to reduced precipitation, below the Turing threshold *P*_*T*_, by forming hexagonal gap patterns. These patterns consist of periodic arrays of circular bare-soil gaps, where any gap is surrounded by six other equidistant gaps. Further decrease in precipitation, below a second threshold, results in a morphology change, where bare soil gaps grow and merge to form patterns of parallel stripes or labyrinthine patterns. Below a third precipitation threshold, a second morphology change takes place, where vegetation stripes breakup to form hexagonal spot patterns. These patterns consist of periodic arrays of circular vegetation spots, where any spot is surrounded by six other equidistant spots.

The common denominator underlying these morphology changes is the increase in bare-soil areas adjacent to vegetation patches, as precipitation is reduced; from bare-soil gaps to bare-soil stripes in the first morphology change, and from bare-soil stripes to bare-soil areas surrounding vegetation spots in the second morphology change. Increasing bare-soil areas compensate for the reduced precipitation by providing an extra source of water – water transport to adjacent vegetation patches through overland water flow, soil-water diffusion, or water conduction by laterally extended roots. These processes act to retain the amount of water available to vegetation patches and buffer the impact of decreased precipitation. As a consequence, and like the one-dimensional case discussed earlier, community structure is expected to remain largely unaffected.

### Insight III: Multiplicity of stable community states and ecosystem management

The multitude of stable hybrid states, i.e. patterned domains of different sizes in otherwise uniform vegetation, open up novel directions for sustainable management of stressed ecosystems that integrate the need for provisioning ecosystem services with the need to preserve species diversity, as explained below.

A spatially uniform community state at high precipitation responds to a drier climate by shifting the community composition to stress-tolerant (high *χ*) species and reducing its functional richness (see Fig. 5a,b). Ecosystem services, such as feeding livestock by grazing, impose further stress and are likely to result in further reduction of functional richness. However, a sufficiently drier climate also induces a Turing instability to periodic patterns and a multitude of stable hybrid states. These states have higher functional richness than those of the uniform and patterned states, separately, and can be even higher (*FR ≈* 0.33) than the functional richness of the original uniform community state before the shift to a drier climate has occurred (*FR* = 0.29, see Fig. 5a). This is because uniform domains give rise to stronger competition and higher water stress, and thus form niches for stress-tolerant species, whereas patterned domains benefit from overland water flow from bare-soil patches, which weakens the competition, and provides niches to fast-growing species.

These results can be used to reconcile the conflicting needs for ecosystem services and preservation of functional diversity, through the management of provisioning ecosystems services by non-uniform biomass removal, so as to induce the formation of hybrid states (Fig. 1c). Moreover, the multitude of stable hybrid states allow to control the relative abundance of fast growing vs. stress tolerant species, and thus the functional evenness, *FE*, of the community; small patterned domains in uniform vegetation (Fig. 7a) or small uniform domains in patterned vegetation (Fig. 7c) give rise to low functional evenness (*FE ≈* 0.67), while domains of comparable sizes give rise to higher evenness (*FE ≈* 0.70).

## Concluding remarks

In this work we studied the interplay between spatial self-organization and community reassembly, using a spatial model of dryland plant communities. This model captures a particular pattern-forming feedback, associated with differential infiltration and overland water flow (Fig. 2), but we expect our findings to hold also for models that capture the feedbacks associated with soil-water diffusion, and water conduction by laterally spread roots, since they all lead to the same bifurcation structure (Fig. 3a). We are not aware of empirical studies of dryland ecosystems that address this interplay, and thus of data that can be used to test our theoretical predictions, but the three insights described above can serve as solid hypotheses for new long-term empirical studies.

Spatial self-organization and community reassembly are different pathways by which dryland ecosystems relax an imposed water stress. The former is a population-level process that involves monospecific plant mortality by *intra*specific competition, and results in an increase of water availability and stress relaxation for the remaining plants. The latter is a community-level process that involves *inter*specific competition and stress relaxation through the displacement of fast-growing species by stress-tolerant species. The relative roles of the two pathways in the overall response to an environmental stress, and the factors that control them, are open questions of direct relevance to the management of ecosystems at risk in climates that become drier.

Spatial self-organization is not limited to dryland ecosystems. Periodic and non-periodic vegetation patterns have been observed and studied in a variety of other ecological contexts including non-drylands plant communities with negative plant-soil feedbacks, such as self-toxicity (***Bonanomi et al., 2014***; ***Marasco et al., 2014***), hydric peat bogs (***Weltzin et al., 2000***; ***Eppinga et al., 2008***), seagrass meadows (***Ruiz-Reynés et al., 2017***), salt marshes (***Zhao et al., 2019, 2021***), aquatic macrophytes in stream ecosystems (***Cornacchia et al., 2018***), and others (***Rietkerk and van de Koppel, 2008***). In most of these systems spatial self-organization intermingles with community dynamics; the findings of this work may be relevant to these contexts as well, or motivate new studies along similar lines.

## Acknowledgments

The research leading to these results has received funding from the Israel Science Foundation under Grant No. 1053/17. During this research BKB has been supported by a PBC postdoctoral fellowship, and JJRB has been supported by a Kreitman postdoctoral fellowship.

## References

Bastiaansen R, Jaïbi O, Deblauwe V, Eppinga MB, Siteur K, Siero E, Mermoz S, Bouvet A, Doelman A, Rietkerk M. Multistability of model and real dryland ecosystems through spatial self-organization. Proceedings of the National Academy of Sciences. 2018; 115(44):11256–11261. doi: https://doi.org/10.1073/pnas.1804771115.

Bertness MD, Callaway R. Positive interactions in communities. Trends in Ecology and Evolution. 1994; 9(5):191–193. doi: https://doi.org/10.1016/0169-5347(94)90088-4.

Bonanomi G, Incerti G, Stinca A, Cartenì F, Giannino F, Mazzoleni S. Ring formation in clonal plants. Community Ecology. 2014; 15:77–86. doi: https://doi.org/10.1556/ComEc.15.2014.1.8.

Borgogno F, D’Odorico P, Laio F, Ridolfi L. Mathematical models of vegetation pattern formation in ecohydrology. Reviews of Geophysics. 2009; 47:RG1005. doi: https://doi.org/10.1029/2007RG000256.

Cornacchia L, van de Koppel J, van der Wal D, Wharton G, Puijalon S, Bouma TJ. Landscapes of facilitation: how self-organized patchiness of aquatic macrophytes promotes diversity in streams. Ecology. 2018; 99:832–847. doi: https://doi.org/10.1002/ecy.2177.

Dawes J, Williams J. Localised pattern formation in a model for dryland vegetation. Journal of mathematical biology. 2015; p. 1–28. doi: https://doi.org/10.1007/s00285-015-0937-5.

De Deyn GB, Raaijmakers CE, Van der Putten WH. Plant community development is affected by nutrients and soil biota. Journal of Ecology. 2004; 92(5):824–834. doi: https://doi.org/10.1111/j.0022-0477.2004.00924.x.

Deblauwe V, Barbier N, Couteron P, Lejeune O, Bogaert J. The global biogeography of semi-arid periodic vegetation patterns. Global Ecology and Biogeography. 2008; 17:715–723. doi: https://doi.org/10.1111/j.1466-8238.2008.00413.x.

Deblauwe V, Couteron P, Bogaert J, Barbier N. Determinants and dynamics of banded vegetation pattern migration in arid climates. Ecological Monographs. 2012; 82:3–21. doi: https://doi.org/10.1890/11-0362.1.

Deblauwe V, Couteron P, Lejeune O, Bogaert J, Barbier N. Environmental modulation of self-organized periodic vegetation patterns in Sudan. Ecography. 2010; 33:1–13. doi: https://doi.org/10.1111/j.1600-0587.2010.06694.x.

DeMalach N, Kigel J, Sternberg M. The soil seed bank can buffer long-term compositional changes in annual plant communities. Journal of Ecology. 2021; doi: https://doi.org/10.1111/1365-2745.13555.

Diaz S, Cabido M. Vive la différence: plant functional diversity matters to ecosystem processes. Trends in Ecology and Evolution. 2001; 16(11):646 – 655. doi: https://doi.org/10.1016/S0169-5347(01)02283-2.

Doedel E. AUTO: A program for the automatic bifurcation analysis of autonomous systems. Congressus Numerantium. 1981; 30:265–284.

Duraiappah AK, Naeem S. Ecosystems and Human Well-being: Biodiversity Synthesis. World Resources Institute, Washington, DC.; 2005.

Eppinga MB, Rietkerk M, Borren Wea. Regular surface patterning of peatlands: Confronting theory with field data. Ecosystems. 2008; 11:520–536. doi: https://doi.org/10.1007/s10021-008-9138-z.

Falik O, Reides P, Gersani M, Novoplansky A. Self/non-self discrimination in roots. Journal of Ecology. 2003; 91:525–531. doi: https://doi.org/10.1046/j.1365-2745.2003.00795.x.

Gilad E, Von Hardenberg J, Provenzale A, Shachak M, Meron E. Ecosystem Engineers: From Pattern Formation to Habitat Creation. Physical Review Letters. 2004; 93(098105). doi: https://doi.org/10.1103/PhysRevLett.93.098105.

Gowda K, Riecke H, Silber M. Transitions between patterned states in vegetation models for semiarid ecosystems. Phys Rev E. 2014 Feb; 89:022701. doi: https://doi.org/10.1103/PhysRevE.89.022701.

Gratani L. Plant Phenotypic Plasticity in Response to Environmental Factors. Advances in Botany. 2014; 2014:208747. doi: https://doi.org/10.1155/2014/208747.

Grünzweig JM, et al. Emergent pathways of ecosystem functioning in a drier and warmer world. In review. 2021;.

von Hardenberg J, Meron E, Shachak M, Zarmi Y. Diversity of Vegitation Patterns and Desertification. Physical Review Letters. 2001; 89(198101). doi: https://doi.org/10.1103/PhysRevLett.87.198101.

Harrison S, Spasojevic MJ, Li D. Climate and plant community diversity in space and time. Proceedings of the National Academy of Sciences. 2020; 117(9):4464–4470. doi: https://doi.org/10.1016/j.jtbi.2012.01.005.

Inderjit, Duke SO. Ecophysiological aspects of allelopathy. Planta. 2003; 217:529–539. doi: https://doi.org/10.1007/s00425-003-1054-z.

Jaïbi O, Doelman A, Chirilus-Bruckner M, Meron E. The existence of localized vegetation patterns in a systematically reduced model for dryland vegetation. Physica D: Nonlinear Phenomena. 2020; 412:132637. doi: https://doi.org/10.1016/j.physd.2020.132637.

Knobloch E. Spatial Localization in Dissipative Systems. Annual Review of Condensed Matter Physics. 2015; 6(1):325–359. doi: https://doi.org/10.1146/annurev-conmatphys-031214-014514.

Lefever R, Lejeune O. On the origin of tiger bush. Bull Math Biol. 1997; 59:263–294. doi: https://doi.org/10.1016/S0092-8240(96)00072-9.

Lejeune O, Tlidi M, Lefever R. Vegetation spots and stripes: dissipative structures in arid landscapes. Int J Quantum Chem. 2004; 98:261–271. doi: https://doi.org/10.1002/qua.10878.

Marasco A, Iuorio A, Cartenì F, Bonanomi G, Tartakovsky DM, Mazzoleni S, Giannino F. Vegetation Pattern Formation Due to Interactions Between Water Availability and Toxicity in Plant–Soil Feedback. Bull Math Biol. 2014; 76:2866—-2883. doi: https://doi.org/10.1007/s11538-014-0036-6.

Mason NWH, Mouillot D, Lee WG, Wilson JB. Functional richness, functional evenness and functional divergence: the primary components of functional diversity. Oikos. 2005; 111(1):112–118. doi: https://doi.org/10.1111/j.0030-1299.2005.13886.x.

Meron E. Nonlinear Physics of Ecosystems. CRC Press, Taylor & Francis Group; 2015. doi: https://doi.org/10.1201/b18360.

Meron E. Pattern formation – A missing link in the study of ecosystem response to environmental changes. Mathematical Biosciences. 2016; 271:1–18. doi: https://doi.org/10.1016/j.mbs.2015.10.015.

Meron E. From Patterns to Function in Living Systems: Dryland Ecosystems as a Case Study. Annual Review of Condensed Matter Physics. 2018; 9:79–103. doi: https://doi.org/10.1146/annurev-conmatphys-033117-053959.

Meron E. Vegetation pattern formation: The mechanisms behind the forms. Physics Today. 2019; 72:30–36. doi: https://doi.org/10.1063/PT.3.4340.

Nathan J, Osem Y, Shachak M, Meron E. Linking functional diversity to resource availability and disturbance: a mechanistic approach for water limited plant communities. Journal of Ecology. 2016; 104:419–429. doi: https://doi.org/10.1111/1365-2745.12525.

Noy-Meir I. Stability of Grazing Systems: An Application of Predator-Prey Graphs. Journal of Ecology. 1975; 63:459–481. doi: https://doi.org/10.2307/2258730.

O’Sullivan JD, Knell RJ, Rossberg AG. Metacommunity-scale biodiversity regulation and the self-organised emergence of macroecological patterns. Ecology Letters. 2019; 22(9):1428–1438. doi: https://doi.org/10.1111/ele.13294.

Parra-Rivas P, Fernandez-Oto C. Formation of localized states in dryland vegetation: Bifurcation structure and stability. Phys Rev E. 2020 May; 101:052214. doi: https://doi.org/10.1103/PhysRevE.101.052214.

Paz-Kagan T, DeMalach N, Zaady E, Shachak M. Resource redistribution effects on annual plant communities in a runoff harvesting system in dryland. Journal of Arid Environments. 2019; 171:103984. doi: https://doi.org/10.1016/j.jaridenv.2019.05.012.

Pielou EC. The measurement of diversity in different types of biological collections. Journal of Theoretical Biology. 1966; 13:131–144. doi: https://doi.org/10.1016/0022-5193(66)90013-0.

van der Putten WH, Bardgett RD, Bever JD, Bezemer TM, Casper BB, Fukami T, Kardol P, Klironomos JN, Kulmatiski A, Schweitzer JA, Suding KN, Van de Voorde TFJ, Wardle DA. Plant–soil feedbacks: the past, the present and future challenges. Journal of Ecology. 2013; 101(2):265–276. doi: https://doi.org/10.1111/1365-2745.12054.

Pérez-Ramos IM, Matías L, Gómez-Aparicio L, et al. Functional traits and phenotypic plasticity modulate species coexistence across contrasting climatic conditions. Nat Commun. 2019; 10:2555. doi: https://doi.org/10.1038/s41467-019-10453-0.

Rietkerk M, Boerlijst MC, van Langevelde F, HilleRisLambers R, van de Koppel J, Kumar L, Prins HHT, De Roos AM. Self-organization of vegetation in arid ecosystems. Am Nat. 2002; 160:524–530. doi: https://doi.org/10.1086/342078.

Rietkerk M, Dekker SC, de Ruiter PC, van de Koppel J. Self-Organized Patchiness and Catastrophic Shifts in Ecosystems. Science. 2004; 305:1926–1929.

Rietkerk M, van de Koppel J. Regular pattern formation in real ecosystems. Trends in Ecology and evolution. 2008; 23(3):169–175. doi: https://doi.org/10.1016/j.tree.2007.10.013.

Ruiz-Reynés D, Gomila D, Tomàs Sintes T, Hernández-García E, Marbà N, Duarte CM. Fairy circle landscapes under the sea. Science Advances. 2017; 3:e1603262. doi: https://doi.org/10.1126/sciadv.1603262.

Sherratt JA. Pattern Solutions of the Klausmeier Model for Banded Vegetation in Semiarid Environments V: The Transition from Patterns to Desert. SIAM Journal on Applied Mathematics. 2013; 73(4):1347–1367. https://www.jstor.org/stable/24510684.

Siteur K, Siero E, Eppinga MB, Rademacher JDM, Doelman A, Rietkerk M. Beyond Turing: The response of patterned ecosystems to environmental change. Ecological Complexity. 2014; 20(0):81–96. doi: https://doi.org/10.1016/j.ecocom.2014.09.002.

Tlidi M, Lefever R, Vladimirov A. On Vegetation Clustering, Localized Bare Soil Spots and Fairy Circles. In: Dissipative Solutions: From Optics to Biology and Medicine, vol. 751 of Lecture Notes in Physics Springer; 2008..

Tzuk O, Ujjwal SR, Fernandez-Oto C, Seifan M, Meron E. Period doubling as an indicator for ecosystem sensitivity to climate extremes. Scientific Reports. 2019; 9:19577. doi: https://doi.org/10.1038/s41598-019-56080-z.

Valentin C, d’Herbes JM, Poesen J. Soil and water components of banded vegetation patterns. Catena. 1999; 37:1–24. doi: https://doi.org/10.1016/S0341-8162(99)00053-3.

Vandermeer J, Yitbarek S. Self-organized spatial pattern determines biodiversity in spatial competition. Journal of Theoretical Biology. 2012; 300:48–56. doi: https://doi.org/10.1016/j.jtbi.2012.01.005.

Weiher E, Freund D, Bunton T, Stefanski A, Lee T, Bentivenga S. Advances, challenges and a developing synthesis of ecological community assembly theory. Philosophical Transactions of the Royal Society B: Biological Sciences. 2011; 366(1576):2403–2413. doi: https://doi.org/10.1098/rstb.2011.0056.

Weltzin JF, Pastor J, Harth C, Bridgham SD, Updegraff K, Chapin CT. Response of bog and fen plant communities to warming and water-table manipulations. Ecology. 2000; 81(12):3464–3478. doi: https://doi.org/10.1890/0012-9658(2000)081[3464:ROBAFP]2.0.CO;2.

Yizhaq H, Gilad E, Meron E. Banded Vegetation: Biological Productivity and Resilience. Physica A. 2005; 356:139. doi: https://doi.org/10.1016/j.physa.2005.05.026.

Yizhaq H, Shachak M, Meron E. A model study of terraced riverbeds as novel ecosystems. Scientific Reports. 2020; 10:3782. doi: https://doi.org/10.1038/s41598-020-60706-y.

Zelnik YR, Kinast S, Yizhaq H, Bel G, Meron E. Regime shifts in models of dryland vegetation. Philosophical Transactions R Soc A. 2013; 371:20120358. doi: https://doi.org/10.1098/rsta.2012.0358.

Zelnik YR, Meron E, Bel G. Gradual regime shifts in fairy circles. Proceedings of the National Academy of Sciences. 2015; 112(40):12327–12331. doi: https://doi.org/10.1073/pnas.1504289112.

Zhao LX, Xu C, Ge ZM, van de Koppel J, Liu QX. The shaping role of self-organization: linking vegetation patterning, plant traits and ecosystem functioning. Proceedings of the Royal Society B: Biological Sciences. 2019; 286(1900):20182859. doi: https://doi.org/10.1098/rspb.2018.2859.

Zhao LX, Zhang K, Siteur K, Li XZ, Liu QX, van de Koppel J. Fairy circles reveal the resilience of self-organized salt marshes. Science Advances. 2021; 7(6). doi: https://doi.org/10.1126/sciadv.abe1100.

